# Virtual colony count study of the inoculum effect of HNP1 against *Staphylococcus aureus* ATCC 29213

**DOI:** 10.64898/2026.04.09.717392

**Authors:** Bryan Ericksen

## Abstract

**Background:** Virtual colony count is a kinetic, 96-well turbidimetric assay that has been used since 2003 to determine the antimicrobial activity of antimicrobial peptides including the defensin HNP1. Virtual colony count results differed from traditional colony counting results in studies of the antimicrobial activity of the human cathelicidin LL-37 and related peptides. The difference could possibly have been caused by an inoculum effect.

**Methods:** The virtual colony count assay was conducted using inocula that varied from 1250 to 1×10^8^ virtual colony forming units (CFUv) per milliliter.

**Results:** The virtual colony count assay demonstrated a pronounced inoculum effect of HNP1 against *Staphylococcus aureus* ATCC 29213, accompanied by biofilm formation observed in the wells of the 96 well plates at all inocula. The *S. aureus* inoculum effect was not as drastic as previously reported for *Escherichia coli*.

**Conclusions:** The inoculum effect is further evidence that biofilm formation is a resistance mechanism used by a variety of bacteria against antimicrobial peptides such as HNP1.

## Introduction

*Staphylococcus aureus* ATCC 29213 has been studied repeatedly as the representative Gram positive strain in a series of publications since 2005 using the virtual colony count (VCC) antibacterial assay. (Ericksen 2005, Pazgier 2012, Rajabi 2012, Wei 2009, Wei 2010, Wu 2005, Wu 2007, Xie 2005a, Xie 2005b, Zhao 2012, Zhao 2013, Zou 2008) In all of these studies, HNP1 was used as the positive control peptide. Therefore, a large body of work exists with this combination of bacterial strain and antimicrobial peptide. In 2013 a side-by-side comparison of VCC and traditional colony count results was published, detailing the activity of the antimicrobial peptide LL-37 and related derivatives against several bacteria including *S. aureus* ATCC 29213. (Pazgier 2013) In that study, survival demonstrated by the traditional colony count method was less than virtual survival demonstrated by VCC for all peptides and bacterial strains tested. This difference could have been due to an inoculum effect. The traditional colony count inoculum was 2.5×10^4^ virtual colony forming units (CFUv)/mL, whereas the standard inoculum of 5×10^5^ CFUv/mL was used for the VCC assays. The inoculum was normalized across the different strains used by turbidity rather than colony forming units (CFU), as explained in the original VCC publication (Ericksen 2005); thus, CFUv was reported rather than CFU. This twenty-fold difference in inoculum, if responsible for the differences in measured activity, would have been the reverse of other inoculum effects reported in the literature (Brook 1989), because the higher inoculum resulted in greater apparent activity.

A modified VCC procedure was used to study the inoculum effect of HNP1 against several strains of bacteria. A study of the inoculum effect of HNP1 against the standard Gram-negative strain used in the history of VCC assays, *Escherichia coli* ATCC 25922, was published in 2020, demonstrating a pronounced inoculum effect at high inocula that almost abrogated activity. (Ericksen 2020) Similarly, the inoculum effect of HNP1 was measured against *S. aureus* ATCC 29213. The results are significant not only from the perspective of the long history of the use of this strain as the Gram-positive representative in VCC assays, but also in the further elucidation of the mechanisms of antimicrobial peptide activity and the mechanisms by which bacteria resist such activity.

## Methods

The VCC procedure was modified for various inocula as previously described (Ericksen 2020), including the use of a centrifuge to concentrate cells for the highest, 1×10^8^ CFUv/mL inoculum, and the application of a strip of Parafilm to the edge of the 96-well plate to retard evaporation and allow all 96 well to be used for the experimental portion of the assay. The change in optical density at 650 nm (ΔOD_650_) was monitored kinetically and the time taken to reach a threshold ΔOD_650_ of 0.02 was calculated using the Visual Basic macro VCC_Calculate (Ericksen 2020). These threshold times were entered into complex Microsoft Excel spreadsheets to calculate virtual survival and virtual lethal doses as described in detail. (Ericksen 2020)

## Results

The inoculum effect of HNP1 against *S. aureus* ATCC 29213 (Figure 1 and Table 1) showed many of the same characteristics as the inoculum effect of HNP1 against *E. coli* ATCC 25922. There was little or no difference in activity at the lower two inocula, 1250 and 2.5×10^4^CFUv/mL, compared to the standard inoculum of 5×10^5^ CFUv/mL. A noticeable inoculum effect was observed at 1×10^7^ CFUv/mL, and a pronounced inoculum effect was observed at the highest inoculum of 1×10^8^ CFUv/mL. Interestingly, at all inocula across a wide range of peptide concentrations, biofilms were observed in the wells of the 96-well plate. (Liao 2025) This result contrasted with *E. coli*, which produced visible biofilms only at the high inocula of 1×10^7^ CFUv/mL or higher. This aspect of the *S. aureus* ATCC 29213 data agrees with the history of the use of this strain in VCC assays where HNP1 was the positive control peptide, when similar biofilms were present at the standard inoculum. There was another difference between *S. aureus* and *E. coli. S. aureus* growth curves (ΔOD_650_ vs. time) deviated from parallel compared to the calibration controls that were not exposed to any antimicrobial agent. (Figure 2) Apparently, HNP1 activity is not completely abrogated by the addition of twice-concentrated Mueller Hinton Broth (2XMHB), indicating the present of salt-tolerant HNP1 activity against *S. aureus* ATCC 29213 but not *E. coli* ATCC 25922 or any of the other strains repeatedly studied in the history of VCC assays such as *Bacillus cereus* ATCC 10876 and *Enterobacter aerogenes* ATCC 13048.

**Table 1.** Virtual lethal doses of triplicate experiments of HNP1 assayed against various inocula of *S. aureus* ATCC 29213 as determined using the VCC assay. Mean values ± standard error of the mean are reported. (1) The three vLD_99.9_ values were 32 < vLD_99.9_ < 64, 14.14, and 8 < vLD_99.9_ < 16.

| | Virtual Lethal Doses ($\mu\text{g/mL}$) | | | |
| --- | --- | --- | --- | --- |
|  | vLD<sub>50</sub> | vLD<sub>90</sub> | vLD<sub>99</sub> | vLD<sub>99.9</sub> |
| 1x10<sup>8</sup> CFUv/mL | 12.85 ± 4.92 | 29.67 ± 9.54 | 64.99 ± 22.83 | 71.87 ± 21.31 |
| 1x10<sup>7</sup> CFUv/mL | 1.89 ± 0.73 | 7.32 ± 2.02 | 20.91 ± 6.44 | 35.89 ± 12.02 |
| 5x10<sup>5</sup> CFUv/mL | 1.74 ± 0.80 | 3.57 ± 1.56 | 11.77 ± 4.55 | 23.98 ± 9.15 |
| 2.5x10<sup>4</sup> CFUv/mL | 1.88 ± 0.95 | 4.14 ± 2.41 | 12.81 ± 5.67 | 23.77 ± 10.33 |
| 1250 CFUv/mL | 2.23 ± 1.22 | 3.37 ± 1.65 | 13.25 ± 4.71 | (1) |

**Figure 1.**
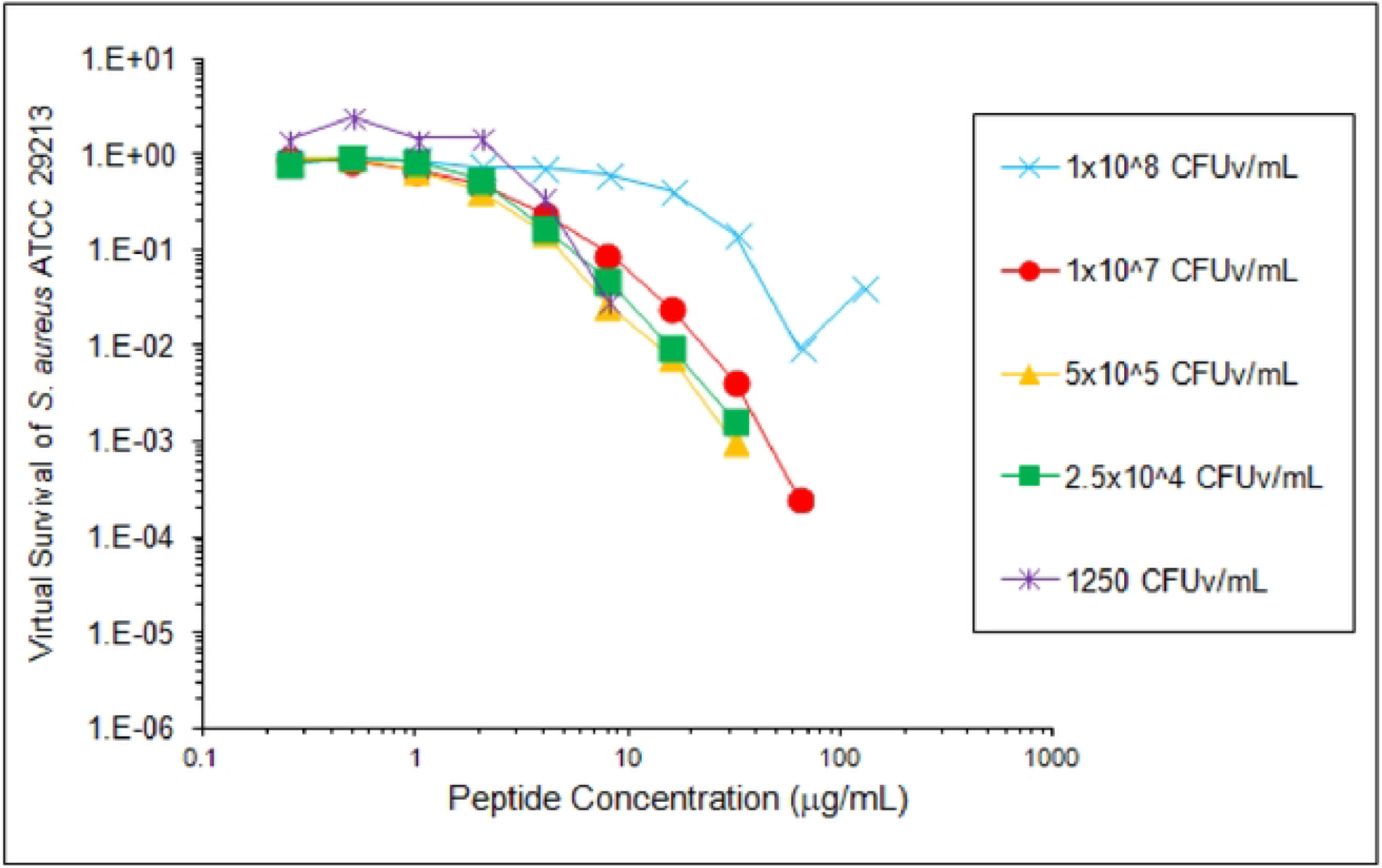
Inoculum effect of HNP1 against *S. aureus* ATCC 29213 as determined by the virtual colony count antimicrobial assay. Virtual survival values were calculated by comparing the threshold times of growth curves of 200 μl batch cultures exposed to a 2-fold dilution series of HNP1 compared to control cultures of the same inoculum exposed to no defensin. Each virtual survival curve is the mean of triplicate experiments. Virtual survival values of zero could not be plotted on a logarithmic scale. Underlying data are available from the Figshare Repository (Ericksen 2026).

**Figure 2.**
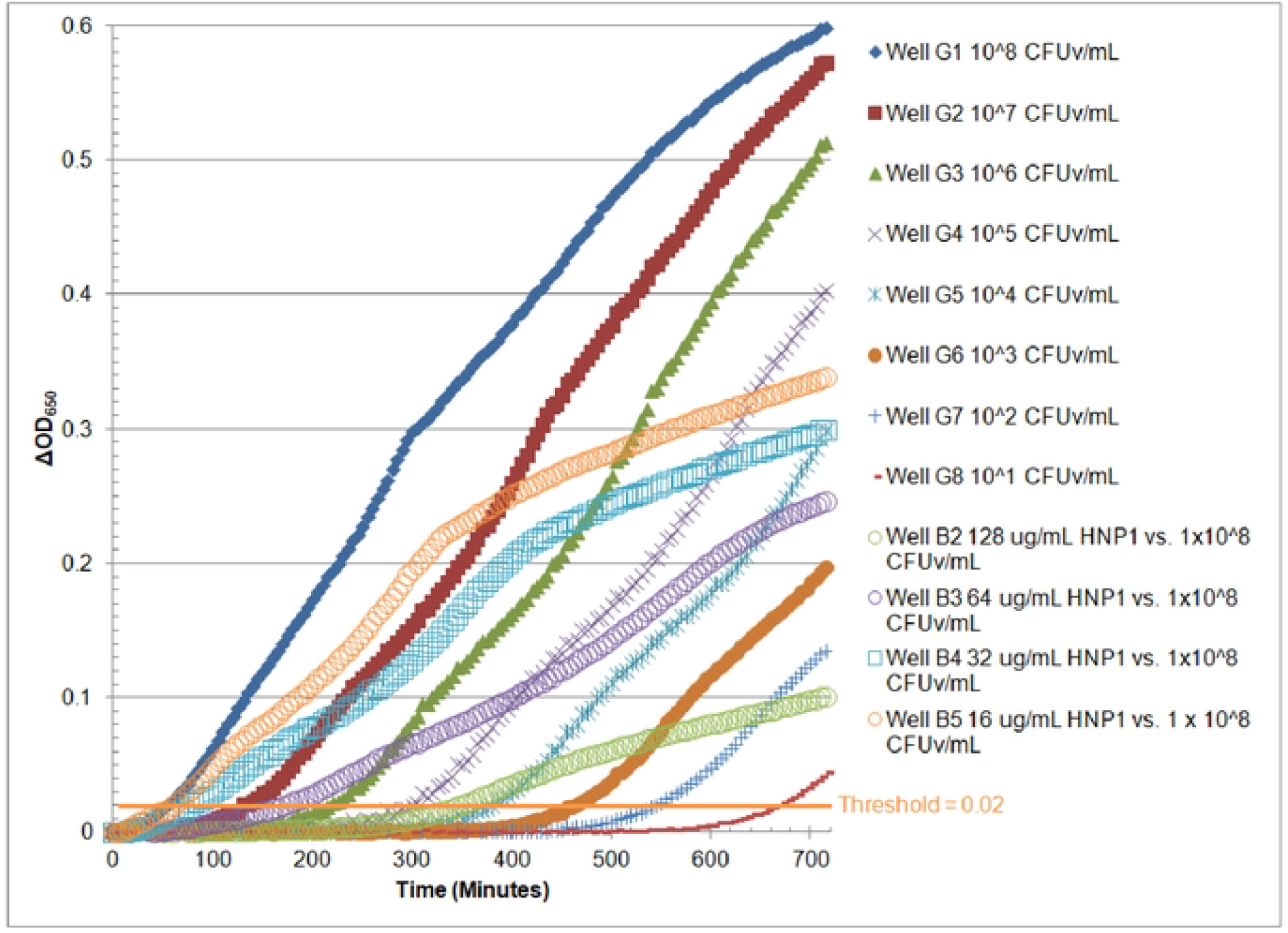
Growth curves of a VCC experiment. Solid markers are a 10-fold calibration dilution of *S. aureus* ATCC 29213 cells not exposed to defensin. Open circles are growth curves of cells exposed to 128 μg/ml (green), 64 μg/ml (lavender), 32 μg/ml (cyan), or 16 μg/ml (orange) HNP1. Growth curves were also markedly non-parallel to the calibration curves at these four HNP1 concentrations at all other inocula tested, with greater deviation the lower the inoculum (data not shown). Minimizing this deviation necessitated choosing a threshold as low as possible while still clearing noise in the initial VCC publication (Ericksen 2005).

## Conclusions/Discussion

The inoculum effect of HNP1 against *S. aureus* ATCC 29213 reinforces other studies of biofilm formation by *S. aureus* (Liao 2025), demonstrating that HNP1 promotes the formation of biofilms that are, in turn, resistant to HNP1 activity. The inoculum effect was not quite as pronounced with *S. aureus* ATCC 29213 as with *E. coli* ATCC 25922, in the sense that there was greater activity against *S. aureus* ATCC 29213 at the high inoculum of 1×10^7^CFUv/mL and virtual survival of less than 1×10^-1^at both 64 and 128 µg/mL at the highest inoculum against *S. aureus* ATCC 29213. However, the presence of *S. aureus* ATCC 29213 biofilms was ubiquitous at many defensin concentrations and all inocula studied, indicating that biofilm formation is a strategy employed by this strain to resist HNP1 activity regardless of cell density. These results may explain why *S. aureus* infections are persistent in spite of the activity of human neutrophils. However, the finding that HNP1 activity was not completely abrogated by the addition of 2XMHB indicates that HNP1 is more effective against *S. aureus* in the presence of physiological salt concentrations than it is against other bacterial strains commonly studied. The bad news for the host regarding biofilm formation may be partially ameliorated by the good news that HNP1 can slow growth rates, possibly indicating that HNP1 has a role in preventing the spread of *S. aureus* colonization to other parts of the body.

The inoculum effect against both *E. coli* and *S. aureus* is consistent with the need for a minimum HNP1 concentration in membranes to be active. This interpretation of the inoculum effect agrees with that of other authors who have discovered inoculum effects of antimicrobial peptides. The first inoculum effect of a defensin was observed in 1993, when the activity of Rabbit NP-1 was tested against *Cryptococcus neoformans*. (Alcouloumre 1993) There was greater minimum inhibitory concentration (MIC) activity observed at the inoculum of 10^4^ cells/mL compared to 10^5^ or 10^6^. Not all studies that varied inoculum size of antimicrobial peptides resulted in an inoculum effect. The activity of omiganan was not affected by inoculum size from 10^3^-10^7^CFU in an ex vivo pig skin colonization model. (Rubinchik 2009) An inoculum effect was demonstrated for Caseicin A and B against *E. coli* when comparing inocula of 10^1^ and 10^3^ cells/mL vs. a standard inoculum of 10^6^ cells/mL. (McDonnell 2011) An inoculum effect using inocula from the neighborhood of the standard inoculum of 5×10^5^ cells/mL to 6×10^6^ cells/mL was also demonstrated for LL-37 against *E. coli*, with increasing MIC as a function of cell density. (Snoussi 2018) A large study of 13 antimicrobial peptides covering more than seven orders of magnitude in inoculated cell density indicated that the inoculum effect is a universal property for the activity of antimicrobial peptides, irrespective of their mechanism of action (Loffredo 2021). It should be noted that HNP1 could not be studied with MIC methods, because the presence of rich media contains sufficiently high salt concentration to abrogate the activity of HNP1, except for the non-parallel VCC growth curves with *S. aureus* ATCC 29213, as noted in the Results section above, which would not be detectable in an overnight MIC assay. It is also noteworthy that a paradoxical data point resulted from the exposure of 128 µg/mL HNP1 to the highest inoculum of 1×10^8^ CFUv/mL. This apparent decrease in activity at the highest defensin concentration could be caused by cell autoaggregation and biofilm formation in that batch culture. Although this phenomenon was observed only in one experiment at the highest cell concentration in this dataset, it occurred at other inocula in the *E. coli* dataset. (Ericksen 2020)

The mechanistic basis for the inoculum effect may involve resistance mechanisms that are more prevalent at high cell concentrations and are part of the process of biofilm formation. For example, bacterial biofilms contain extracellular polymeric substances (EPS) including nucleic acids and polysaccharides. (Ikuma 2013) EPS may contain polymers such as capsular polysaccharides and RNA that bind and inhibit defensins. (Ericksen 2015) However, biofilms were observed at all cell concentrations in the VCC studies of *S. aureus* ATCC 29213, indicating that this resistance mechanism is important for both dilute, predominantly planktonic cells and higher cell concentrations.

Regarding the original impetus to conduct the study of the inoculum effect, it is apparent that the difference in activity between VCC and traditional colony count observed in the 2013 *Biochemistry* study (Pazgier 2013) cannot be explained by an inoculum effect. The simplest explanation of the difference is that VCC threshold time delays can be caused both by bactericidal killing and by the lag phase preceding exponential growth. Comparing calibration threshold times with traditional colony count results, lag times of approximately 60-90 minutes were observed. Apparently, after exposure to antimicrobial peptides such as LL-37, the survivors require some time to repair or replace damaged membranes, cell walls, or intracellular targets before entering into exponential growth. Further studies are warranted to elucidate the mechanistic underpinnings of the inoculum effect.

## Data Availability

Data underlying Figure 1 are available from the Figshare Repository (Ericksen 2026).

## Acknowledgments

I thank Wuyuan Lu for providing antimicrobial peptides and helpful discussions.w

